# SAM-based Automatic Workflow for Histology Cyst Segmentation in Autosomal Dominant Polycystic Kidney Disease

**DOI:** 10.1101/2024.09.02.610807

**Authors:** Pablo Delgado-Rodriguez, Roman Kinakh, Rafael Aldabe, Arrate Munoz-Barrutia

## Abstract

Autosomal Dominant Polycystic Kidney Disease (ADPKD) is a genetic disorder characterized by the development of numerous cysts in the kidneys, ultimately leading to significant structural alterations and renal failure. Detailed investigations of this disease frequently utilize histological analyses of kidney sections across various stages of ADPKD progression. In this paper, we introduce an automated workflow leveraging the Segment Anything Model (SAM) neural network, complemented by a series of post-processing steps, to autonomously segment cysts in histological images. This approach eliminates the need for manual annotations or preliminary training phases and enables precise quantification of cystic changes over entire kidney sections. Application of this method to sequential histology images across the development timeline of ADPKD in mice demonstrated a notable increase in the proportion of diseased tissue from 8 to 12 weeks and from 12 to 16 weeks, with the cysts appearing progressively lighter. Our workflow not only surpasses the performance of the existing Cystanalyser tool but also offers enhanced flexibility and accuracy in full-image segmentation. The developed workflow is made publicly accessible to facilitate its adoption as an efficient tool for rapid and reliable cyst segmentation in histological studies.

## Background

Autosomal Dominant Polycystic Kidney Disease (ADPKD) represents the most common form of hereditary renal disorder, for which there is currently no cure [1]. Characterized by the progressive enlargement of kidney structures -primarily within the collecting ducts and distal nephron segments [2] -ADPKD leads to the formation of fluid-filled cysts that significantly alter the anatomy and impair the function of the kidneys, with potentially fatal outcomes. ADPKD is often not detected in humans until its advanced stages. The current gold standard for its radiological diagnosis is renal ultrasonography, an inexpensive and non-invasive method of examination [3]. Other techniques like Magnetic Resonance Imaging (MRI) are also used in diagnosis [4]. Although the disease can manifest from an early age, screening is generally recommended for children if there is a parental history of ADPKD.

To elucidate the pathogenic mechanisms of ADPKD, researchers employ various analytical approaches, including both human and animal studies. Animal models, particularly genetically modified mice, provide a controlled setting to precisely monitor disease progression and derive insights at specific developmental stages. These models facilitate the visual examination of cyst proliferation through sequential imaging, enabling quantitative analyses of cystic areas.

Among the imaging modalities utilized, histology offers exceptionally high resolution, allowing for detailed observation of individual cysts within kidney tissues [5, 6, 7, 8]. Longitudinal studies in ADPKD mouse models typically involve the euthanasia of mice at predetermined intervals post-induction, followed by the extraction and histological processing of their kidneys. This process yields a series of high-resolution 2D images at each time point, within which cyst segmentation and area quantification can be performed. Manual segmentation, however, is labor-intensive and repetitive.

Recent advances in deep learning have significantly enhanced the capabilities of image segmentation algorithms, traditionally requiring extensive training or fine-tuning on specific datasets. Although such models can provide very detailed results in histological images [9], they need the generation of manually annotated datasets for model optimization, a process often at odds with the goal of minimizing manual labor.

Addressing this challenge, we utilize the Segment Anything Model (SAM) [10], a network designed to perform initial cyst segmentation that is subsequently refined through automated post-processing, yielding final cyst masks from histological images. The adaptability of this model allows it to recognize objects across a wide range of image types without bespoke training, presenting a robust solution for automated segmentation.

The application of SAM for histological image processing has been explored recently. In [11], SAM is applied to different scale histological images, achieving the segmentation of tumors, cell nuclei and other structures. This process is performed using input prompts to guide the algorithm and assist in correct segmentation. In [12], an improved version of SAM is lightly retrained to be able to process histological images more accurately, including also prompting for segmentation production, and focusing on tumor segmentation. Our approach differs from these studies in that we aim to process whole large-scale images that include complete kidney sections, and segment small structures (cysts) appearing along the whole sample. Moreover, our method does not require any training whatsoever, making it suitable for direct and easy application to quantitatively evaluate ADPKD evolution on kidneys.

In this manuscript, we present an automatic segmentation methodology that obviates the need for pre-training, thereby streamlining the process of cyst segmentation in histological images of kidneys from ADPKD-affected mice. We detail the procedural steps, discuss the results from a study tracking kidney development at 2, 4, 8, 12, and 16 weeks post-ADPKD induction, and compare the efficacy of our method against the Cystanalyser approach [13].

## Implementation

### Image acquisition

The animal models used in this study were Pax8rtTA [14], TetO-Cre [15], and PKD2^fl/fl^ mice, sourced from the Baltimore PKD Research and Clinical Core Center. These models were specifically chosen for the conditional inducible knockout (KO) of the polycystic kidney disease 2 (PKD2) gene, facilitating the induction of ADPKD via controlled doxycycline administration.

Breeding and housing of the mice were conducted at the CIMA Universidad de Navarra animal facilities, and all the procedures were approved by the Universidad de Navarra Animal Research Ethics Committee (081c-19). Starting at four weeks of age, the mice were administered doxycycline-laced drinking water continuously for a two-week period to induce the disease. Post-induction, mice were sequentially sacrificed at predefined time points - 2, 4, 8, 12, and 16 weeks - to track the progression of the disease. The kidneys were then harvested, sectioned, and stained with Hematoxylin and Eosin (H&E). These sections were subsequently scanned using an Aperio CS2 slide scanner (Leica Biosystems), producing high-resolution histological images.

Each kidney yielded ten distinct histological images, each representing a different section of the organ. The resolution of these images was maintained at 2.008 μm per pixel, although the total image sizes varied. In some cases, kidneys were imaged together with liver cuts to increase efficiency and as much data as possible from each image acquisition. In these images, the liver cuts were later manually removed from the image for the current study, since the focus of our method was on kidney cysts. Figure 1 provides a schematic of the imaging protocol over the different time points, alongside example images from each stage.

**Figure 1.**
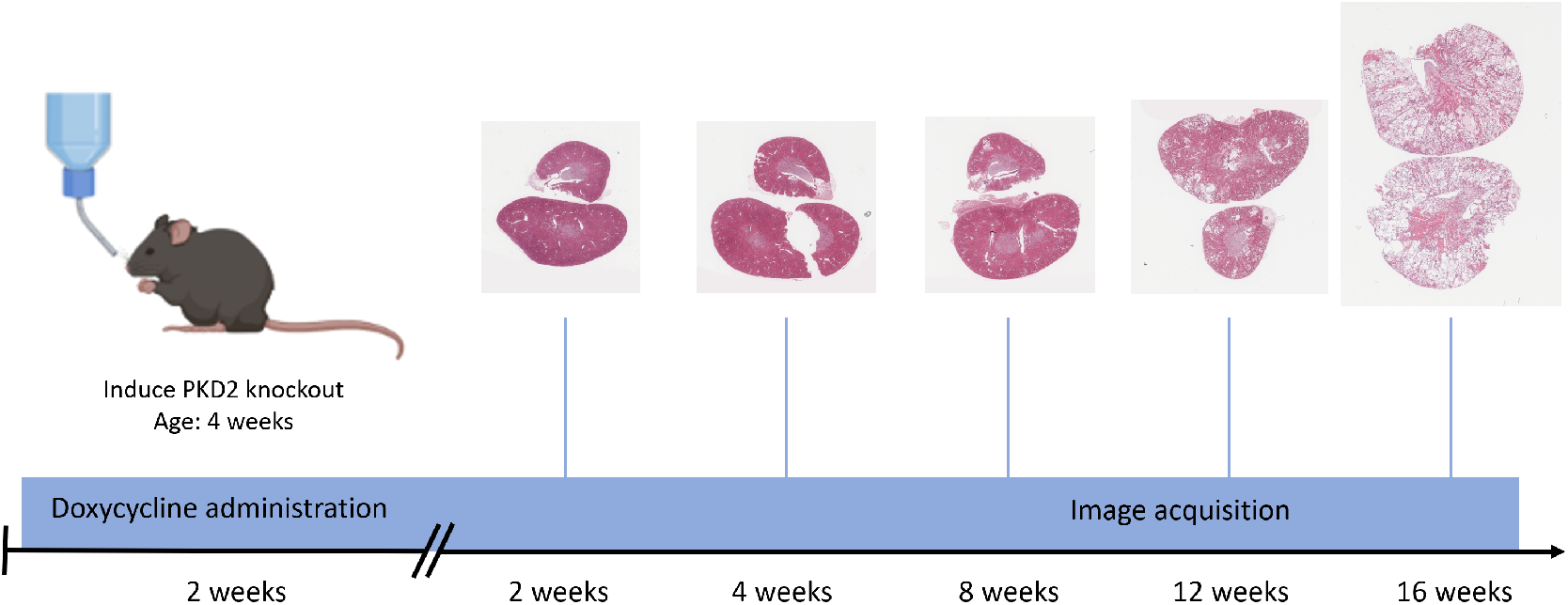
Schematic overview of the image acquisition process. Mice were induced with ADPKD via doxycycline administration for 2 weeks, followed by sequential sacrifices at specified intervals. Histological images of their kidneys were captured at each time point. Representative images of pathological kidneys are included in the figure.

The aggregate data regarding the number of pathological and control samples analyzed at each time point are detailed in Table 1.

**Table 1:** Number of samples at different time points.

| Time (weeks) | 2 | 4 | 8 | 12 | 16 |
| --- | --- | --- | --- | --- | --- |
| Pathological kidneys | 4 (40 images) | 3 (30 images) | 4 (40 images) | 4 (40 images) | 4 (40 images) |
| Control kidneys | 0 | 0 | 0 | 6 (60 images) | 0 |

### Automatic segmentation workflow

We developed a fully automated segmentation workflow that utilizes the pre-trained Segment Anything Model (SAM), followed by several post-processing steps to enhance the accuracy and usability of the segmented outputs. The entire process is detailed as follows:

1. **Patch Division:** Initially, each histology image is divided into square patches. The side length of each patch is determined to be one-fifth of the image’s horizontal length.
2. **Preliminary Cyst Detection:** The SAM algorithm is applied to each patch individually, identifying cysts and other structures as distinct objects.
3. **Patch Reassembly:** Following detection, the patches are reassembled to form a cohesive global mask of the original image.
4. **Size Filtering:** Objects disproportionately large relative to their respective patch size are eliminated to avoid false positives.
5. **Border Correction:** The peripheral pixels of each patch, predominantly marked as background, are adjusted by extending the intensities of nearby detected objects to these borders.
6. **Patch Integration:** Neighboring patches are analyzed to ensure that cysts divided during the initial patch creation are seamlessly merged, maintaining uniform intensity.
7. **Edge Analysis:** Any object with a high proportion of straight edges relative to its bounding box perimeter is discarded, as these are likely artifacts from patch borders or corners.
8. **Intensity Homogeneity:** The standard deviation of grayscale intensities within each object is calculated. Objects with a deviation exceeding a predetermined threshold are removed to retain only homogeneously appearing elements (cysts).
9. **Kidney Masking:** Otsu’s thresholding method is applied to a grayscale version of the original image to facilitate the identification of up to four major kidney regions. These are chosen between the largest connected components with axis dimensions exceeding one-sixth of the image width. The convex hull of these regions is then computed to create an approximate mask of the entire kidney.
10. **Final Mask Application:** The comprehensive kidney mask is combined with the cyst mask, ensuring that only cysts within the kidney boundaries are retained. The resulting mask shows a different intensity for each of the cysts found in the image, and the background as 0.

Figure 2 presents a summarized overview of the segmentation process, illustrating both the entire image and a zoomed-in section to highlight detailed segmentation effects.

**Figure 2.**
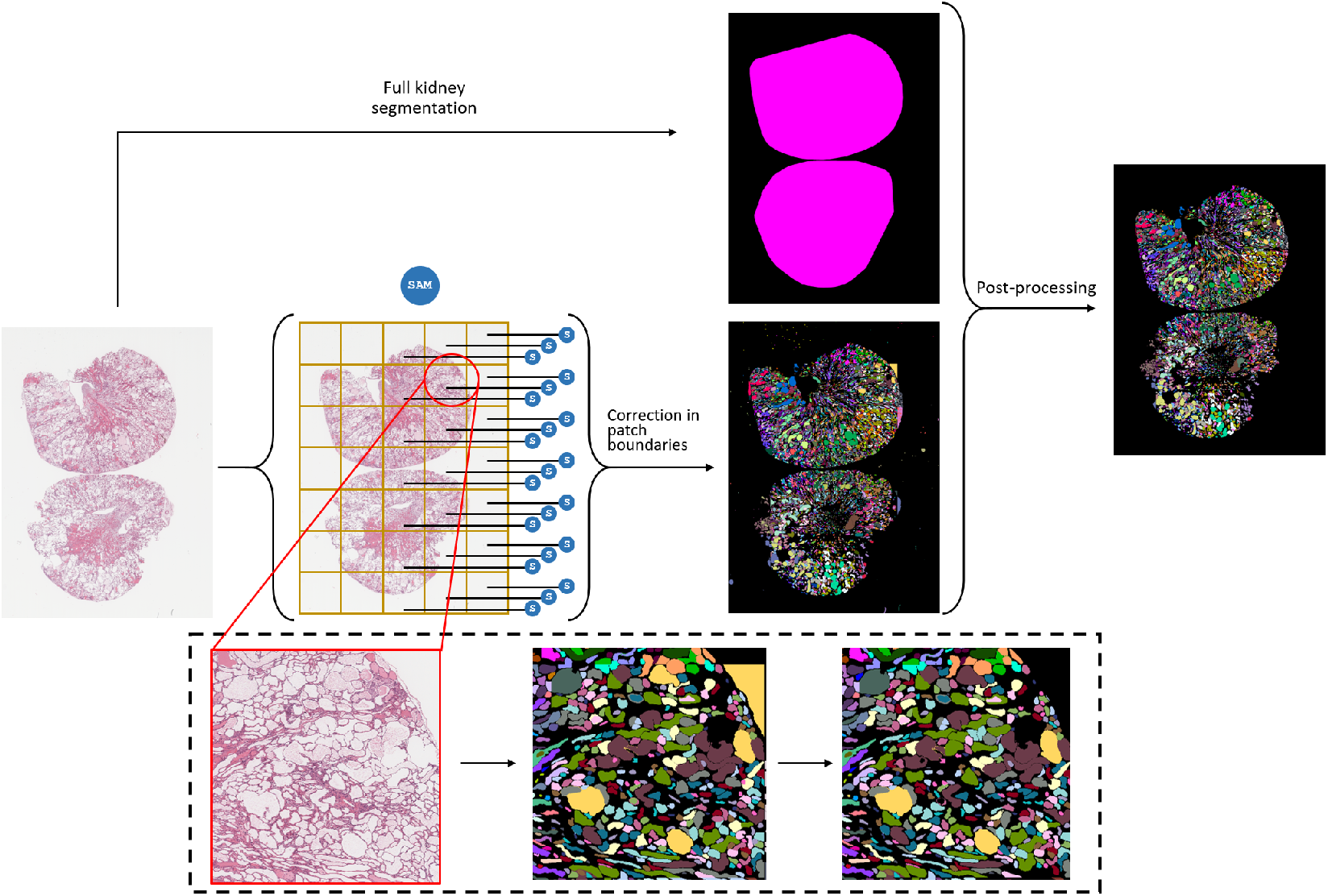
Schematic representation of the cyst segmentation workflow. Initial patch-based segmentation is enhanced through systematic post-processing, culminating in the integration with a kidney-specific mask to delineate the final cyst regions accurately.

This segmentation workflow was implemented using a Nvidia GeForce RTX 3090 GPU with 24 GB of memory, which resulted in significantly faster processing when applying the deep learning model than CPU-based methods. The version of SAM employed is accessible at https://github.com/facebookresearch/segment-anything.

### Quantification of ADPKD development

Following the creation of the final cyst masks, a series of quantitative measures were employed to evaluate the progression of ADPKD and to assess the efficacy of our segmentation workflow. The following metrics were calculated for each different kidney, composed by 10 images. The mean and standard deviations of each metric were obtained between all the cuts of each specific kidney:

- **Proportion of Cystic Area:** The total area occupied by cysts in the image was quantified as a proportion of the overall kidney area, extracted from the full kidney segmentation mask. This ratio provides a direct measure of disease progression.
- **Number of Cysts per mm**^2^: While the previous variable gave a quantification of the area affected by ADPKD, this one allows to evaluate the speed of new cysts formation, more than their increase in size. Both quantities, combined, can offer an overall view of the kidney’s transformation during this process
- **Mean Cystic Area:** The average area of the individual cysts was computed, offering insights into the growth patterns of each individual cyst over time.
- **Mean Cystic Eccentricity:** This measure was used to evaluate the elongation of the cysts, where a value closer to zero indicates a perfect circle, and values closer to one suggest an ellipse. This was computed to check whether there was a clear trend in cyst shape depending on the disease’s stage.

To statistically analyze these data, one-way Analysis of Variance (ANOVA) tests were conducted to compare the means of each set of measurements. This included comparisons between the mean values from all kidneys at a time point vs all kidneys from the consecutive one to detect significant changes over the course of the disease, as well as comparisons between pathological and control kidneys at the 12-week mark to assess disease impact.

Although the control kidneys should not show any cysts, the SAM workflow, being an automatic method without retraining, also detects some gaps that are present within the organ and are not produced by the disease. This can be seen as missegmentations; however, most times these structures have proven difficult to distinguish visually from cysts even for an expert, and other methods that have been tested, like Cystanalyser, also mark them as cysts. Despite these additional segmentations, the initial detections in control kidneys can be considered as a baseline over which cyst segmentations are observed. The evolution in cyst formation and any derived metrics are going to be shown nonetheless as a function of time in the pathological data, and these increases showcase the behavior of ADPKD despite the initial stage containing non-cystic structures. Given this, the comparison with control kidneys at 12 weeks has been included to assess the differences between a regular kidney and a kidney affected by a later-stage ADPKD.

The combined use of these metrics provides a comprehensive overview of ADPKD progression in the murine models, facilitating a robust evaluation of the segmentation methodology implemented.

### Comparison with Cystanalyser

To assess the performance of our segmentation workflow relative to existing tools, we conducted a comparative analysis with Cystanalyser [13]. Our approach was to apply Cystanalyser to each image in the most automatic way possible, in order to analyze the differences between the two automatic methods, without any specific input required from the user. Initial attempts to apply Cystanalyser to full image scans were unsuccessful due to the limitations of the software in processing large images. Consequently, we adopted a patch-based approach for this comparison. A series of smaller patches (600 x 600 pixels) were selectively extracted from various histological sections, ensuring they accurately represented the interior of the kidney. A single kidney cut was chosen for each different stage of disease progression, including a control specimen at 12 weeks. Two patches were extracted from each of these cuts, so a total of 12 patches were evaluated.

The Cystanalyser software’s interface required the input of each parameter to be performed in a specific order, ensuring they had been set adequately. It should be noted that steps such as the need to set a specific pixel size before applying any method were not well documented. Failing to perform this action at the beginning resulted in the program closing suddenly. This and other details posed some difficulties when attempting to apply it in the most automatic way possible. This interface allows for a wide range of cyst radii settings, but for our purposes, this range was set to include any possible structure. This translates into a range from 3 to 1000 pixels (6 to 2000 *μ*m). The ‘Kidney’ segmentation option was chosen before clicking the “Automatic Cyst Recognition” button. For comparison, the regions corresponding to these same patches were extracted from the whole image cyst mask previously generated by our SAM-based workflow.

Manual segmentations of cysts found within the patches were produced to evaluate both methods against them, creating a Ground Truth (GT) for performance comparison. Gap regions of a large-enough size were marked as cysts within these ground truth segmentations, including the control kidneys. This was done to account for the recognition of these empty regions, that in later stages of the disease cannot be visually distinguished from the cysts, and form a baseline over which to observe the additional formation of cysts during ADPKD.

To check both methods against the manual segmentations, each GT mask was compared with the corresponding prediction masks of both Cystanalyser and the SAM workflow. In these comparisons, for each cyst *A* detected in the prediction mask, for every intensity *B* of the GT mask that overlapped with *A*, the Jaccard similarity index *J* (Equation 1) was computed between them. The *B* with the highest Jaccard index was kept as a matching object for that value of *A*.

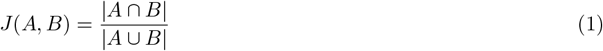

To assign a matching pair *A* - *B*, the condition shown in Equation 2 also needed to be met, to ensure the cyst was recognized and the overlap was high enough. This condition is derived from the Segmentation Evaluation Protocol of the Cell Tracking Challenge [16]

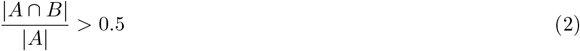

These metrics allowed to account for the number of correctly detected (matching) and incorrectly detected (not matching) cysts for each prediction mask coming from Cystanalyser or the SAM workflow. The correctly detected cysts ratio, *CCR* (number of correctly detected cysts over the total number of detected cysts), was then obtained as a measure of performance for each image comparison (Equation 3, where *N*_*corr*_ is the number of correctly detected cysts and *N*_*inc*_ the number of incorrectly detected cysts).

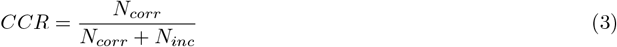

Total cystic area proportion, *CAP*, was also analyzed (total area of the cysts over area of the whole patch). Absolute differences of this quantity (*dif CAP* ) were calculated between the GT’s *CAP* (*CAP*_*GT*_ ) and each prediction’s *CAP* (*CAP*_*pred*_) as seen in Equation 4

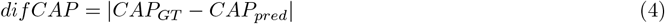

This comparison between two different methods aims to highlight the adaptability and accuracy of our SAM-based workflow against the established Cystanalyser software as an automatic alternative to it. The measures taken allowed to tackle both the correct detection of the individual cysts and the whole cystic area.

## Results and Discussion

### PKD evolution measurements

The progression of ADPKD is quantitatively analyzed through a series of metrics derived from the segmented images. In all corresponding figures, distinct annotations indicate statistically significant differences in mean (one-way ANOVA, 95% confidence); different letters represent significant changes between consecutive time points, while an asterisk marks significant discrepancies between pathological and control specimens at the 12-week interval.

- **Cystic Area Proportion:** Figure 3A illustrates the ratio of the area occupied by cysts to the total kidney area, providing an estimation of the kidney’s degradation at various stages of ADPKD. Initially, this ratio remains relatively stable through weeks 2, 4, and 8. A marked increase is observed at week 12 and continues to rise into week 16, indicating a rapid progression of the disease, which substantially deforms the kidney structure and impairs its function. This trend confirms the progressive nature of ADPKD, validating the efficacy of our segmentation approach for rapid and reliable disease monitoring.
- **Mean number of cysts per mm**^2^: The evolution of the mean number of cysts per mm^2^ can be seen in Figure 3D. These numbers are very similar at 2 and 4 weeks, with a certain increase at 8 weeks, but not significant enough. At 12 weeks, there is, however, a very marked and statistically significant increase in cysts per mm^2^. From 12 to 16 there is again a significant increase, which in turn follows a very similar trend than the total cystic area. These results support the observations of the previous graph, with the first three time points showing an initial development phase with slow cyst growth, while from 8 to 12 weeks, ADPKD causes a rapid formation of multiple cysts that continues up to the 16-week point.
- **Mean Area per Cyst:** As depicted in Figure 3C, the average area of each cyst (mm^2^) does not show significant changes over time, nor between pathological and control specimens at 12 weeks. These results, together with the numbers shown in Figure 3B, suggest that ADPKD progression is characterized more by an increase in cyst number rather than an enlargement of existing cysts. As the disease spreads, more and more cysts appear and occupy a larger portion of the kidney’s area.
- **Cyst Eccentricity:** The mean eccentricity of cysts, ranging from 0 (perfect circle) to 1 (highly elongated), is shown in Figure 3D. The data indicate that the shape of the cysts remains consistent over time in the pathological kidneys. However, there is a significant difference between the pathological and control mice at 12 weeks. Given that some non-cystic structures are being detected in the control kidneys, as mentioned in the previous section, it can be inferred that cysts seem to exhibit a more elongated shape, on average, than healthy kidney structures such as transversal vessel cuts.

**Figure 3.**
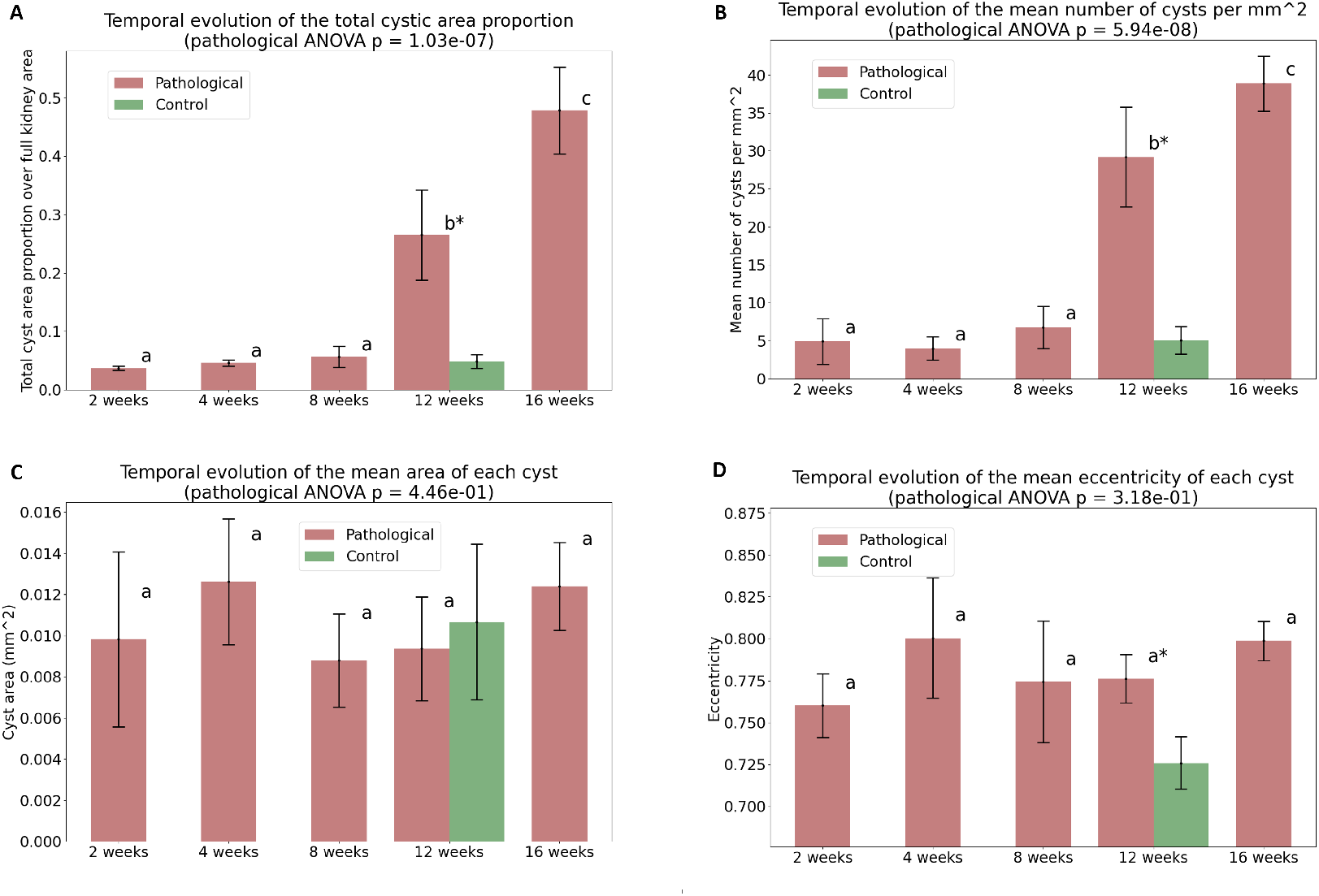
Various graphical representations of ADPKD progression metrics extracted from the SAM workflow’s segmentations. Mean values with standard deviations shown as a bar around these means. Different letters denote significant differences in means between temporally consecutive groups, and asterisks indicate significant differences between pathological and control kidneys (one-way ANOVA, 95% confidence). A global one-way ANOVA at 95% confidence has also been applied for the difference in means of all pathological measurements with its p-value shown within the title of each graph. A: Proportion of the area occupied by cysts relative to the total kidney area; B: Mean number of cysts per mm^2^; C: Mean area of each cyst; D: Mean eccentricity of each cyst.

Overall, we observed a gradual increase in the diseased area up to the 8-week mark, followed by a significant escalation at 12 and 16 weeks. This pattern matched the one shown by the total number of cysts per mm^2^, indicating a critical phase of rapid disease progression at these time steps. Despite these dynamic changes in the cystic area and number, the mean area and eccentricity of the cysts remained consistent, suggesting that the progression in ADPKD primarily involves an increase in cyst number rather than changes in their size or shape.

### Cystanalyser comparison

This section presents the results of a detailed comparison between our SAM-based workflow and the segmentations produced by Cystanalyzer. The mean correctly detected cysts ratio, *CCR*, was computed for both methods between each of the studied patches. The mean difference in total cystic area proportion, *dif CAP* was also obtained for comparison and are shown in Table 2 together with other metrics.

**Table 2:** Comparison of Cystanalyser vs SAM workflow.

|  | Cystanalyser | SAM workflow |
| --- | --- | --- |
| Applicable to whole image | No | **Yes** |
| Can be run fast without a GPU | **Yes** | Not optimally |
| Can extract measures from a complete histology slice | No | **Yes** |
| Mean ± std Correct Cysts Ratio (CCR) | $0.162 \pm 0.157$ | **$0.590 \pm 0.277$** |
| Mean ± std difference in Total Cystic Area Proportion (difCAP) | $0.092 \pm 0.026$ | **$0.019 \pm 0.019$** |

The results show that both methods miss-segment some proportion of the cysts. However, it can be observed how Cystanalyser vastly over-segments too many structures as cysts. While the SAM workflow misses some of the cysts, the resulting differences are not as notable as those of Cystanalyser. The mean Corrected Cysts Ratio (*CCR*) is 0.590 for SAM, which shows most of the detected cysts are correct. On the other hand, it is 0.162 for Cystanalyser, which marks many small light areas as cysts, increasing the incorrect cyst count significantly and thus reducing *CCR*. Similar results can be seen for the mean difference in total cystic area proportion (*dif CAP* ), which is much lower for the SAM workflow, 0.019, than for Cystanalyser, 0.092. This shows that the overall measure of ADPKD spread is much more accurate using our SAM pipeline than Cystanalyser, which introduces too many additional detected cysts that contribute to a significant overestimation of the cystic area. An example of this behavior can be seen in Figure 4, where one of the patches corresponding to the 4-week time point is shown, together with the GT, Cystanalyser, and SAM workflow segmentations. Specific values of *CCR* and *dif CAP* are also shown for both methods on this patch, with the SAM method achieving a high *CCR*_*SAM*_ to 0.725 while Cystanalyser includes many additional cysts, lowering its *CCR*_*cystAn*_ to 0.109. This result is also reflected in the values for *dif CAP*, with a much lower quantity for *dif CAP*_*SAM*_ = 0.020, which explains how that mask is much closer to the original in terms of the estimated diseased area than the mask from Cystanalyser (*dif CAP*_*cystAn*_ = 0.109).

**Figure 4.**
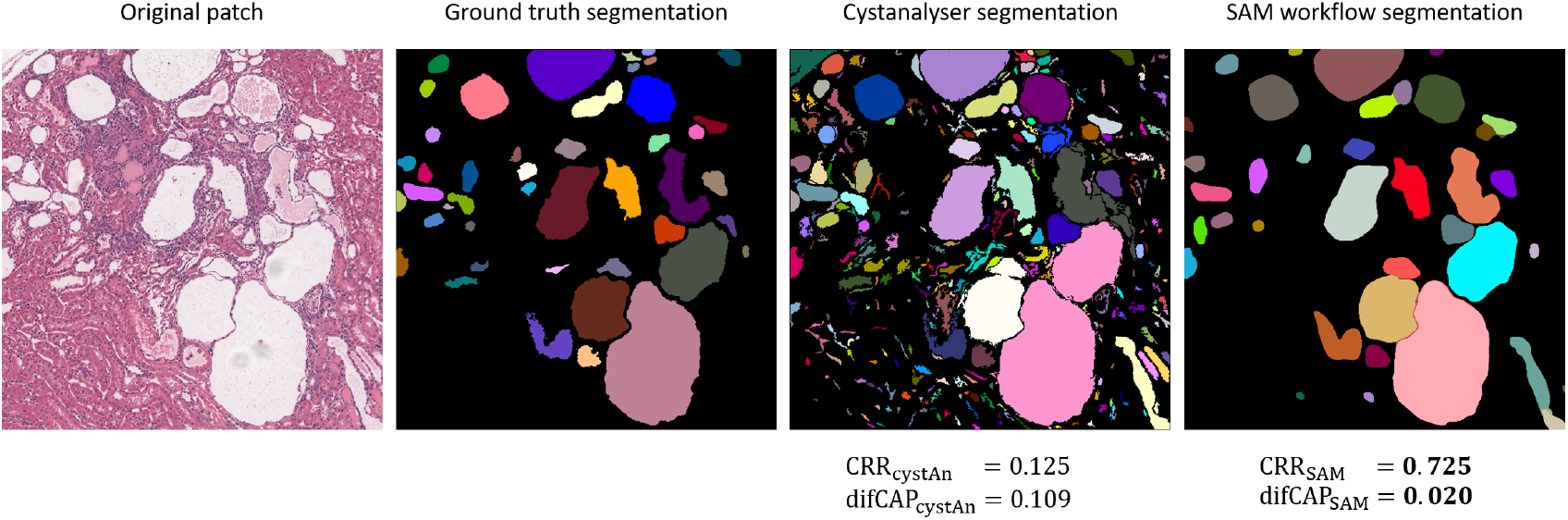
An example of the cyst segmentation masks produced from one of the histology patches. The first image is the original patch. The second image is the ground truth segmentation. The third and fourth images are the Cystanalyzer and SAM workflow segmentation, respectively. Different cysts are marked with different colors. For the two prediction segmentations, their Corrected Cysts Ratio (CCR) and difference in Total Cystic Area Proportion (difCAP)

The ability of our SAM workflow to analyze entire images significantly outperformed the limited patch-based approach of Cystanalyser. This capability provides a more holistic and representative measurement of the entire kidney slice, enhancing the reliability of the disease progression assessment. Besides this, Cystanalyser’s interface requires some specific order of input parameters that is not clearly stated in the documentation, which can be troublesome for the user that intends to apply it in a fast and direct manner. Moreover, its outputs do not include a segmentation mask, which had to be created through an additional script from the produced .xml file with its segmentation details. Offering a mask is a standard output that facilitates the integration of this method with additional processing steps and would be beneficial for Cystanalyser. Our SAM workflow, on the contrary, has a segmentation mask as its primary output.

To summarize, our SAM-based workflow demonstrated superior capabilities to Cystanalyser, avoiding the segmentation of many non-cystic structures detected by the latter. Although our method is not optimized for non-GPU environments, it significantly outperforms Cystanalyser in comprehensive and automated cyst detection and segmentation, with comparisons between both shown in Table 2. The benefits of using our SAM workflow extend beyond mere technical performance, providing a more effective and efficient tool for researchers studying ADPKD’s pathological impacts, that will be able to apply it to whole histological images.

### Limitations and Potential Enhancements

While our study provides significant insights into cyst segmentation using SAM, it also presents some limitations that warrant further investigation;

1. **Generalizability to Human Samples:** The application to mouse models might not fully capture the complexity of human ADPKD. Future research could focus on validating and adapting the workflow for use with human kidney samples.
2. **3D Image Segmentation:** Current segmentation is limited to 2D slices. Extending this method to 3D could offer a more holistic view of cyst distribution and kidney morphology.
3. **Dependency on High-Performance Computing:** The requirement for GPU processing limits the accessibility of our workflow. Developing a more computationally efficient version could broaden its application, especially in settings with limited technological resources.
4. **Segmentation errors:** Our workflow commits some errors, such as identifying, in some cases, glomeruli as cysts. Given the unsupervised nature of the method, it is not possible to directly obtain a perfect solution. Additional post-processing steps could be added in the future to refine the resulting masks as much as possible.
5. **Automated Pathological Feature Recognition:** Integrating the capability to recognize and quantify other pathological changes like fibrosis or inflammation alongside cyst segmentation could enhance the utility of this tool in comprehensive kidney disease analysis.

## Conclusions

In conclusion, our study has successfully developed and validated a novel automatic segmentation workflow utilizing the SAM model for histological images of mouse kidneys affected by ADPKD. This method has proven effective in delineating cystic structures within kidney tissue without the need for manual annotations or any type of training, accommodating the analysis of large-scale histology images to provide comprehensive insights into the disease progression.

Our workflow is available at https://github.com/pdelgado248/SAM workflow histology cysts, providing a straight-forward, effective tool for the quantification of ADPKD effects in histological studies. Researchers can download and implement this method to facilitate their investigations into this complex disease, furthering our understanding and potentially aiding in the development of therapeutic strategies.

Our findings demonstrate that the SAM-based workflow can accurately track the progression of ADPKD, showing a significant increase in the cystic area and number of cysts per mm^2^ during the later stages of the disease. This reflects a critical transition in disease dynamics, characterized by a rapid increase in cyst formation and fluid accumulation. The ability of our workflow to outperform existing tools like Cystanalyzer in terms of segmentation performance and ease of use underscores its potential utility in both research and clinical settings.

Future research will focus on adapting and validating our SAM-based workflow for use with human samples, expanding its capabilities to 3D imaging for comprehensive cyst morphology analysis, correcting miss-segmentations and optimizing computational efficiency for broader accessibility. Additionally, efforts will aim to integrate more pathological features such as fibrosis and inflammation into the analysis and enhance machine learning techniques for better accuracy.

## Ethics Approval

Experimental procedures conducted during this project adhered to the European Communities Council Directive (2010/63/EU) and national rules (RD53/2013, ECC/566/2015) for care of laboratory animals. All protocols received approval from the Ethics Committee for Animal Experimentation from Centro de Investigacion Medica Aplicada (CIMA) Universidad de Navarra.

## Funding

This work was partially funded by Ministerio de Ciencia, Innovación y Universidades through an FPU fellowship (FPU19/02854) and the UC3M fellowship: PIPF programa “Inteligencia Artificial”. Additional funding was provided by Ministerio de Ciencia, Innovación y Universidades, Agencia Estatal de Investigación (MCIN/AEI/10.13039/501100011033/), under Grants PID2023-152631OB-I00, TED2021-129392B-I00 and RTC-2017-6600-1, co-financed by European Regional Development Fund (ERDF), “A way of making Europe”. Partial support also came from Grant 0011-1411-2019-000074 from proyectos de I+D estratégicos (RIS3) from Departamento de Desarrollo Económico del Gobierno de Navarra. This work was further supported by the European Union through the Horizon Europe program (AI4LIFE project, grant agreement 101057970-AI4LIFE). Views and opinions expressed are, however those of the authors only and do not necessarily reflect those of the European Union. Neither the European Union nor the granting authority can be held responsible for them.

## Notes

https://github.com/pdelgado248/SAM_workflow_histology_cysts

### Competing Interest Statement

The authors have declared no competing interest.

### Summary of Updates

This version of the manuscript has been revised to update the Funding section.

